# Invasive plant species interact with drought to shift key functions and families in the native rhizosphere

**DOI:** 10.1101/2023.04.24.538159

**Authors:** Cassandra L. Ettinger, Marina L. LaForgia

## Abstract

Interactions between species invasions and climate change have the potential to drive changes in plant communities more than either factor alone. One pathway through which these effects can occur is via changes to the rhizosphere microbial community. Invasive plants can alter the taxonomic and functional makeup of these microbial communities, which may affect natives’ abilities to compete with invaders. At the same time, climate change is leading to more frequent extreme wet and dry events, shifting the composition of microbial taxa available in the soil. Understanding the response of plant communities to these combined global change drivers requires a comprehensive approach that assesses the relationship between plant competition and belowground rhizosphere microbial community responses. Here we use a field experiment in a California grassland with a set of six native annual forbs (i.e., wildflowers) and three invasive annual grasses to test how competition with invasive plants alters both identity and function in the native rhizosphere microbiome, and whether competition between these groups interacts with rainfall to amplify or ameliorate these microbial shifts. Metagenomics of rhizosphere communities revealed that drought combined with competition from invaders altered a higher number of functions and families in the native rhizosphere compared to invasive competition alone or drought alone. This suggests invasion-driven shifts in the microbial community may be involved in weakening natives’ ability to cope with climate change, especially drought. Understanding the role of the microbial community under invasion and climate change may be critical to mitigating the negative effects of these interacting global change drivers on native communities.

## Introduction

The invasion of novel species can drive a multitude of cascading effects on the structure and function of native communities, but climate change combined with invasion has the potential to amplify and worsen these effects. Invasive plants alter soil chemical and microbial composition, outcompete natives for light, water and nutrient resources, and change the pollinator community, often to the advantage of invaders over native residents (Ehrenfeld, 2010; Reinhart & Callaway, 2006; Rodríguez-Caballero et al., 2020; Traveset & Richardson, 2014). The effects of invasive species, however, inherently depend on the availability of resources - thus, under a rapidly changing climate, the strength of these effects may shift and novel pathways of invasive dominance may arise. One of the least understood pathways contributing to invasive dominance over natives lies in the rhizosphere microbiome. While there is ample evidence that invaders can shift the rhizosphere microbiome, and some evidence suggesting that these invader-induced changes may aid in the dominance of invasives over natives (LaForgia et al., 2022), how climate interacts with invasive species to alter the rhizosphere microbiome remains poorly understood.

The microbial community that inhabits the rhizosphere, the region of soil directly surrounding plant roots, often forms close associations with host plants (Edwards et al., 2015). Members of this community can be detrimental (pathogenic), neutral (commensalistic) or beneficial (mutualistic) to host plant fitness (Thrall et al., 2007). For example, beneficial microbes can help host plants with amelioration of biotic and abiotic stress (Doubková et al., 2012; Yang et al., 2009), acquisition of nutrients (Richardson et al., 2009), promotion of plant growth (Qu et al., 2020), and pathogen resistance (Ritpitakphong et al., 2016). In return, host plants provide benefits to the microbial community, often through the release of sugar compounds into rhizosphere soil; these sugars double as attractants to entice microbes into beneficial interactions (Hartmann et al., 2008; Paterson et al., 2007). As a result, rhizosphere microbial communities tend to be host plant-specific (Aleklett et al., 2015; Dawson et al., 2017) and are critical for the persistence of native plant species (Chung et al., 2015; David et al., 2019). When a native plant community becomes invaded, the addition of novel plant hosts may alter the microbial composition of the rhizosphere, either directly by recruiting a different set of microbes from the bulk soil than natives (Batten et al., 2006) or indirectly through the release of compounds that might select for a different community composition in the bulk soil (Ulbrich et al., 2022), affecting the availability of microbes to neighboring native plants.

We face a daunting knowledge gap about how climate change will interact with microbial communities to affect soil and plant health. In addition to being closely tied to plant host identity, the dynamic makeup of rhizosphere microbial communities is strongly influenced by abiotic soil variables including soil water availability (Fierer et al., 2012). For example, functional and taxonomic microbial diversity tend to be lower in more arid regions (Fierer et al., 2012; Maestre et al., 2015) and microbial activity tends to decrease during drought (Acosta-Martinez et al., 2014; Legay et al., 2017; Ochoa-Hueso et al., 2018). Further, changes in water availability have the capacity to shift the relationship between plants and microbes (Cheng et al., 2019), and complicating matters, both plants and microbes can also shape surrounding abiotic soil conditions. As climate change drives increases in rainfall variability in many systems, especially semi-arid and arid ones, understanding the effects on the rhizosphere microbiome and their plant hosts becomes increasingly important.

In California annual grasslands, rainfall variability mediates the composition of native forbs (i.e., wildflowers or non-grass herbaceous plants) and invasive grasses through time. In general, wet years favor competitively dominant invasive grasses, while successive drought years indirectly favor subdominant natives via decreased invasive grass competition (Dudney et al., 2017; LaForgia et al., 2018). Further, recent work suggests that native species may be resilient to changes in climate in the absence of invasive grasses (LaForgia et al., 2020). In the presence of these invasives, however, the negative effects of drought are intensified and the beneficial effects of increased rainfall are dampened (LaForgia et al., 2020). Invasive grasses in this community also drive shifts in the rhizosphere microbial community composition (Batten et al., 2006) which may be contributing to invasive dominance over natives (LaForgia et al., 2022). In order to understand the effects of invasion on native plant communities under climate change, microbial shifts must be assessed under different levels of rainfall availability at the local level.

Recent technological advances in molecular and computational biology now enable robust surveying of the genetic content of entire communities through metagenomic sequencing. Unlike high-throughput amplicon sequencing, which only enables taxonomic profiling of communities, metagenomic sequencing allows us to use genes to assess both who is in the community and their predicted functional potential. Additionally, new advances in machine learning tools enable us to draw insights about microbial community dynamics from metagenomic data. The application of SourceTracker, a Bayesian classifier, to metagenomic data is effective at predicting the proportion of a community assembled from known sources (Knights et al., 2011; McGhee et al., 2020), enabling better understanding of microbial community assembly under competing host plants. Utilizing amplicon data, these methods predicted that the joint rhizosphere microbiome of invasive and native plant pairs was equally sourced from both the native host and the invasive host in a well-watered greenhouse experiment (LaForgia et al., 2022). Neighbor-induced changes in the rhizosphere microbiome are greatest under strong competition (Ulbrich et al., 2022) and tend to shift towards those of the dominant competitor (Hortal et al., 2017; Lozano et al., 2019). Thus under drought, which has been shown to intensify the negative effects of invasive competition on natives (LaForgia et al., 2020), we might expect to see a higher proportion of microbes sourced from invasive hosts. By using metagenomics in combination with these machine learning approaches, we can more precisely infer the functional impact that invaders have on the native community via the rhizosphere microbial community, as well as gain insights into complicated ecological processes driving these impacts.

To understand how rainfall variability and competition from invasive grasses affect the rhizosphere microbial community of native forbs, we manipulated watering regimes and plant hosts in a field experiment. We then used metagenomics to assess rhizosphere differences between watering and plant host treatments in terms of taxonomy and function as well as host contribution to rhizosphere community assembly. Based on previous findings, we hypothesized that (1) function and taxonomy in the rhizosphere would differ between plant hosts and between watering treatments, but the combined effect of invasive competition and watering treatments on native forb rhizospheres would be larger than invasive grass competition or watering treatments alone. To test this hypothesis we investigated community dissimilarity (i.e., beta diversity) and differential (relative) abundance of key taxa and functions across treatments. (2) We also hypothesized that, while microbial functional and taxonomic communities in invaded native communities under increased watering would be assembled equally from invader and native hosts, as was seen in LaForgia et al (2022), under drought, a majority of the community would be assembled from invaders. To test this hypothesis we implemented a Bayesian classifier to predict the proportion of the microbial community sourced from each host when natives compete with invasives.

## Methods

### Experimental set-up

In November 2017, we set up a field experiment in an annual serpentine grassland at the University of California McLaughlin Natural Reserve in the Inner North Coast Range (N 38°52’, W 122°26’). This reserve has a Mediterranean climate with cool, wet winters and hot, dry summers.

We utilized an existing rainfall manipulation experiment in a lush serpentine (high Mg, low Ca) area of the reserve, which hosts both abundant invasive grasses and high native diversity. The existing experiment consists of 30 watered plots along three watering lines emanating from a rainfall catchment system with each plot centered on a sprinkler that casts water over a 3-m radius (Mini Rotor Drip Emitters, Olson Irrigation) (Figure 1a). Of the 30 watered plots, we randomly chose 10 for the present study. The experiment also utilized 10 drought plots interspersed with watered plots. Drought plots were each covered by a 3 x 3 m shelter constructed following the design of DroughtNet (wp.natsci.colostate.edu/droughtnet) except that the removable roofs intercepted 100% of rainfall. Roofs were placed on the shelters from approximately 1 December 2017 to 1 March 2018. In previous years, these watering treatments led to an average decrease in soil moisture of 42% in the drought plots and an average increase in soil moisture of 27% in the water addition plots (data not published). Watering was applied weekly from December 1 to February 1 to maintain total rainfall at or slightly above its 30-y average. Ten unmanipulated control plots were also interspersed with treatment plots. All plots were no less than 4 meters from each other.

**Figure 1.**
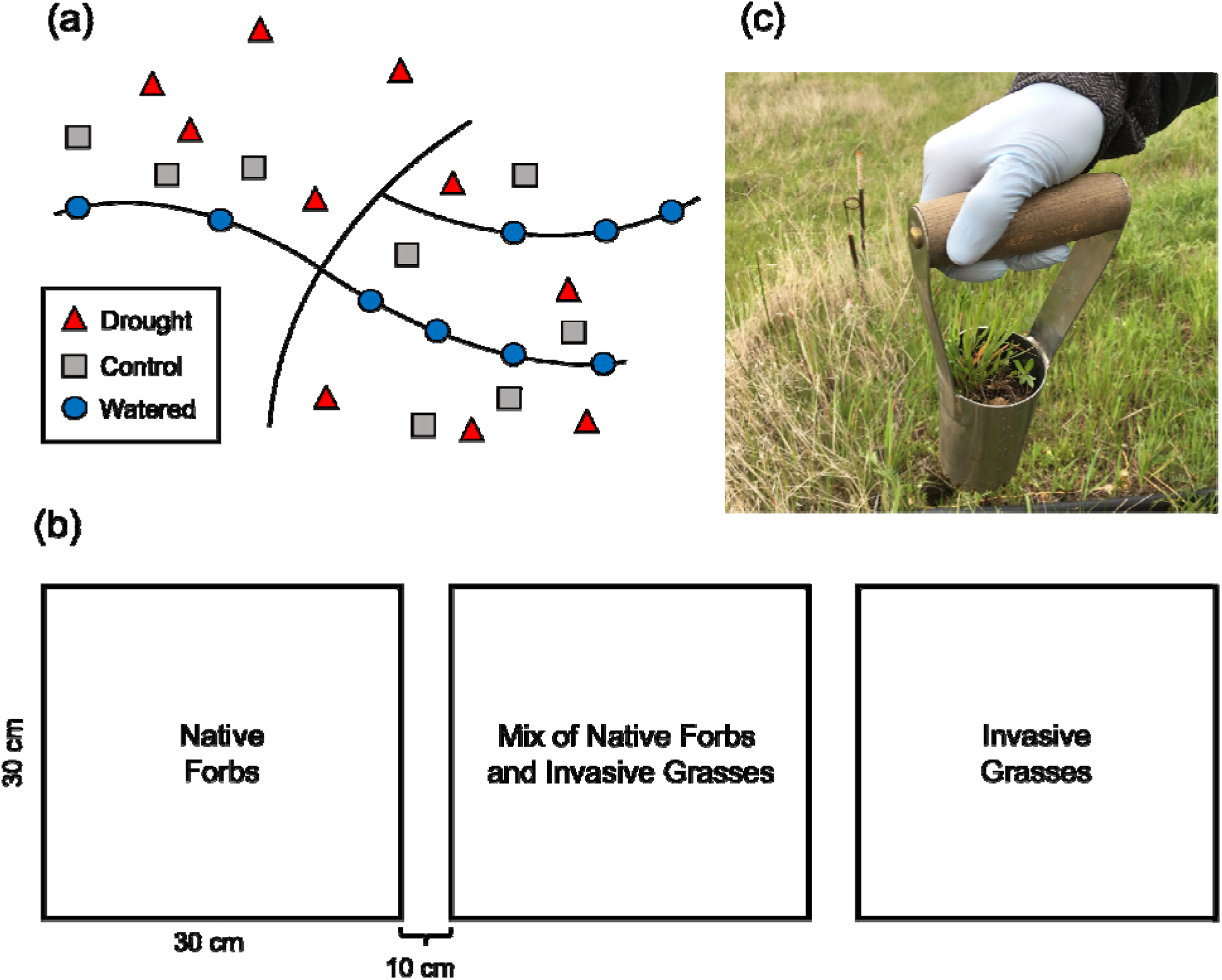
(a) Diagram of plot locations at the field site. Watered plots (blue circles) included a sprinkler along watering lines. Control plots (gray squares) and drought plots (red triangles) were interspersed within the lush serpentine area of the site. (b) Diagram of subplot setup within each plot. (c) Photograph of rhizosphere core sampling method with a native-invasive mix. A sterilized bulb planter was used to core a representative assemblage of plants within each subplot treatment.

Within each of these 30 experimental plots we cleared aboveground vegetation and seeded in three treatment subplots (Figure 1b): (1) invasive grasses, (2) native forbs, and (3) a mixture of invasive grasses and native forbs. We used two different seed mixes of forbs (mix 1: *Lasthenia californica*, *Clarkia purpurea*, and *Agoseris heterophylla*; mix 2: *Calycadenia pauciflora*, *Hemizonia congesta*, and *Plantago erecta*). All seeds were collected from the site during the previous spring/summer and subplots were hand weeded halfway through the study to maintain plant host treatments as seeded.

As these species have vastly different flowering phenologies, and development can alter microbial composition (Chaparro et al., 2014; Xiong et al., 2021; J. Zhang et al., 2018), we chose to sample the rhizosphere prior to any flowering so that all plants were sampled during the same life stage. This occurred after the majority of rainfall had occurred (April 5th, 2018). Plants and associated soil were removed from each subplot using a hand bulb planter (Figure 1c), which allowed us to collect a representative of each species included in each treatment at the same time. Samples were placed into a brand new Ziplock bag and processed immediately. We sterilized the bulb planter between subplots and recorded all species present in the core sample (Table S1). For a step-by-step field protocol see Ettinger and LaForgia (Ettinger & LaForgia, 2021).

### Molecular analysis

Excess soil was shaken gently from the roots associated with each subplot. The collective roots from each subplot, with soil still attached, were then placed in a 50 mL conical tube of autoclaved nanopure water and stirred vigorously to collect the soil rhizosphere fraction associated with the plant community in each subplot similar to rhizosphere collection in Edwards et al. (Edwards et al., 2015). Samples were then transported to UC Davis for processing on ice. Conical tubes were centrifuged at 4000 g for 1 min to obtain rhizosphere soil pellets, excess water was decanted and soil pellets were transferred to new 1.5 mL tubes for storage at −80C until further processing. We chose a subset of 66 samples (out of a total of 150 samples) for DNA extraction and subsequent metagenomic analysis based on final plant community composition within the subplots. We prioritized sequencing of rhizosphere samples that accurately represented our treatments as seeded (i.e., without volunteers from alternative functional groups; Table S1).

Samples were randomized using a random number generator prior to DNA extraction. Approximately ∼250 mg of the rhizosphere soil pellet was used for DNA extractions using the MoBio PowerSoil DNA extraction kit (MO BIO Laboratories, Inc.). Extractions were performed following manufacturer’s instructions, with one modification (samples were bead beaten on the “homogenize” setting for 1 minute during step 5). We additionally extracted DNA from three replicates each of a positive technological control using the ZymoBIOMICS Microbial Community Standard (Zymo Research Inc.), four replicates of a negative technological control from autoclaved nanopure water, and three replicates of a negative technological control where no sample was added during DNA extraction. DNA from replicate controls for each control type were pooled to produce representative control samples and these were included as controls during metagenomic sequencing.

### Data generation

DNA samples were randomly aliquoted into wells in a 96 well plate using a random number generator and sent to the Integrated Microbiome Resource (IMR) at Dalhousie University for metagenomic sequencing on an Illumina NextSeq 550. Briefly, the IMR used one nanogram of each sample for Nextera XT (Illumina) library preparation, per the manufacturer’s instructions except clean-up and normalization which were completed using the Just-a-Plate 96 PCR Purification and Normalization Kit (Charm Biotech). Equal amounts of all barcoded samples were then pooled and sequenced on a shared 150+150 bp PE Illumina NextSeq 550. The resulting libraries produced a total of 625,865,988 reads, with a median of 9,133,838 reads per sample (min: 27,026, max: 16,911,776). The raw sequence reads generated for this metagenomic project were deposited at Genbank under BioProject accession no. <u>PRJNA925931</u> (Table S2).

### Data processing

Quality filtering of sequence reads was performed using bbMap v. 37.68 with the following parameters: qtrimlJ=lJrl, trimqlJ=lJ10, maq=lJ10 (Bushnell, 2022). MEGAHIT v.1.1.3 was then used to co-assemble the filtered reads from all samples into a single metagenome assembly (Li et al., 2015). Prior to downstream analysis, we used the anvi’o v.7 metagenomic workflow to process the metagenomic data (Eren et al., 2015). Briefly, this involved using ‘anvi-gen-contigs-database’ to make a database of contigs from our co-assembly and then predict open reading frames using Prodigal v.2.6.2, resulting in 8,829,098 putative genes (Hyatt et al., 2010). Open reading frames were exported from anvi’o using ‘anvi-get-sequences-for-gene-calls’ and functionally annotated using Eggnog-mapper v. 1.0.3 (Cantalapiedra et al., 2021) and InterProScan v. 5.52-86.0 (Jones et al., 2014). Taxonomy was predicted for each gene using Kaiju v.1.5.0 against the NCBI BLAST nonredundant (nr) protein database, including fungi and microbial eukaryotes, v.2017-05-16 (Menzel et al., 2016). Finally, Salmon v. 1.3.0 was used to quantify the number of reads from each individual sample that mapped to predicted genes (Patro et al., 2017).

Salmon read counts and Kaiju taxonomies for each gene were imported into R for analysis and visualization (R Core Team, 2022). Count tables were filtered to remove genes with no read counts and those not seen more than 3 times in at least 5% of the samples. To identify genes associated with possible contaminant taxa, we used decontam’s (Davis et al., 2018) prevalence method with a threshold of 0.5 to identify genes with a higher prevalence in negative controls (e.g, no-sample technological controls). Using this threshold, we identified 143 contaminant genes which were subsequently removed prior to downstream analysis, resulting in a dataset of 128,022 genes.

Genes taxonomically assigned by Kaiju as being from eukaryotes or which were unable to be classified were further investigated using MMseqs2 v. 13.45111 (Steinegger & Söding, 2017) against the NCBI non-redundant database (downloaded December 13, 2022). We then removed 10,860 genes assigned by both software with a eukaryotic, viral or unclassified taxonomic assignment in order to focus on bacterial and archaeal genes. We then further removed 13,266 genes that did not have an associated COG functional annotation. The final set after filtering totaled 104,065 genes. For downstream beta-diversity analysis, samples were rarefied to a read count of 66,880. This value was chosen after examining rarefaction curves in order to balance maximizing the number of reads per sample while minimizing the number of samples removed from downstream analysis. This resulted in the sample with the lowest read count (35,618) being excluded from further analysis.

To understand how function in the microbial community differed between watering and plant host treatments, we agglomerated gene counts based on shared COG function using tax_glom in the phyloseq package (McMurdie & Holmes, 2013). To understand how the taxonomic composition of the community differed between watering and plant host treatments, we agglomerated gene counts at the genus level also using tax_glom. To assess native and invasive contribution of function and identity to community assembly in the native-invasive rhizosphere, we agglomerated gene counts to both COG function and taxonomic family. This was done on rarefied gene counts for downstream beta-diversity analysis and on raw gene counts for downstream differential abundance analysis and community assembly investigations.

### Beta diversity (community dissimilarity) analyses

To assess how the functional beta diversity of rhizosphere microbiomes varied across watering and plant host treatments, we conducted a PERMANOVA on Bray-Curtis dissimilarity between samples of the rarefied and agglomerated gene function data. This was done using adonis2 in the vegan package with by = margin and 9999 permutations (Oksanen et al., 2022). We also included plot as a factor to account for our nested experimental design. We conducted a similar analysis as above to examine how taxonomic beta diversity in the rhizosphere varied across watering and plant host treatments with the agglomerated genus data. To assess which treatment combinations were driving results, we conducted pairwise adonis contrasts on focal treatment combinations.

Given that plant species composition varied within plant host treatments, we also assessed whether presence or absence of specific host species were drivers of beta diversity within invasive, native or native-invasive mixes. Taking the previously calculated Bray-Curtis dissimilarities, we used the envfit function to fit host species presence/absence onto a principal coordinates analysis ordination made with the cmdscale function, both from the vegan package. This was done for both the agglomerated functional dataset and the agglomerated taxonomy dataset. We tested for correlations between overall plant host composition and Bray-Curtis dissimilarity using Mantel tests with 9999 permutations.

### Functional and taxonomic differential (relative) abundance analyses

To assess how function in the rhizosphere varied across watering and plant host treatments, we used DESeq2 (Jonsson et al., 2016; Love et al., 2014) on the agglomerated and unrarefied gene function data to examine the log_2_fold change (i.e. differential [relative] abundance, hereafter referred to as differential abundance) of each functional category between pairwise contrasts of plant host groups (native forbs, invasive grasses, and native-invasive mixes) and watering treatment (control, watered, drought). We used Benjamini-Hochberg corrections to adjust *p*-values and kept functions that had an adjusted *p*-value <0.05. We then summed log_2_fold changes to the broader COG category to assess overall patterns of functional differences.

We conducted a similar analysis as above to examine how taxonomy in the rhizosphere varied across watering and plant host treatments with the agglomerated taxonomy data, then summed the resulting significant log_2_fold changes to the family level.

### Host plant contribution to assembly of genes in native-invasive mixes

To assess the proportion of the joint microbial community sourced from native forbs or invasive grasses in mixed native-invasive subplots (similar to LaForgia et al (2022)), we employed a Meta-SourceTracker approach using Sourcetracker2 (Knights et al., 2011; McGhee et al., 2020). Sourcetracker2 was used to estimate the fraction of families or functions detected in sinks (mixed subplots) that might originate from sources (invasive grasses, native forbs, or unknown habitats). We used source and sink rarefaction depths of 25,000. We obtained the predicted proportion of each taxonomic family and of each COG function in mixed subplots originating from each source using the --per_sink_feature_assignments option. We used Wilcoxon signed-rank tests to assess differences in the SourceTracker2 predicted proportions between different sources (invasive grasses, native forbs, or unknown) for the microbial communities across different watering regimes.

## Results

### Taxonomic and functional beta diversity differ between watering treatments and plant hosts

As stated above, we hypothesized that function and taxonomy in the rhizosphere would differ between plant hosts and watering treatments, with the interaction between invasive competition and watering treatments showing the largest effects on native forb rhizosphere. Contrary to our hypothesis, the interaction between watering treatment and plant host treatment did not have a significant effect on functional beta diversity (p = 0.213) nor on taxonomic beta diversity (p = 0.336). Both functional beta diversity and taxonomic beta diversity did however significantly differ between watering treatments (function: p < 0.001, Figure 2a; taxonomy: p < 0.001, Figure S1a) and between plant host treatments (function: p = 0.044, Figure 2b; taxonomy: p = 0.011, Figure S1b). For both taxonomy and function, significant differences between plant hosts were found between native forbs and native-invasive mixes (function: p = 0.024; taxonomy: 0.009), while differences between watering treatments were significant between all treatment combinations (see Table S3 for full results).

**Figure 2.**
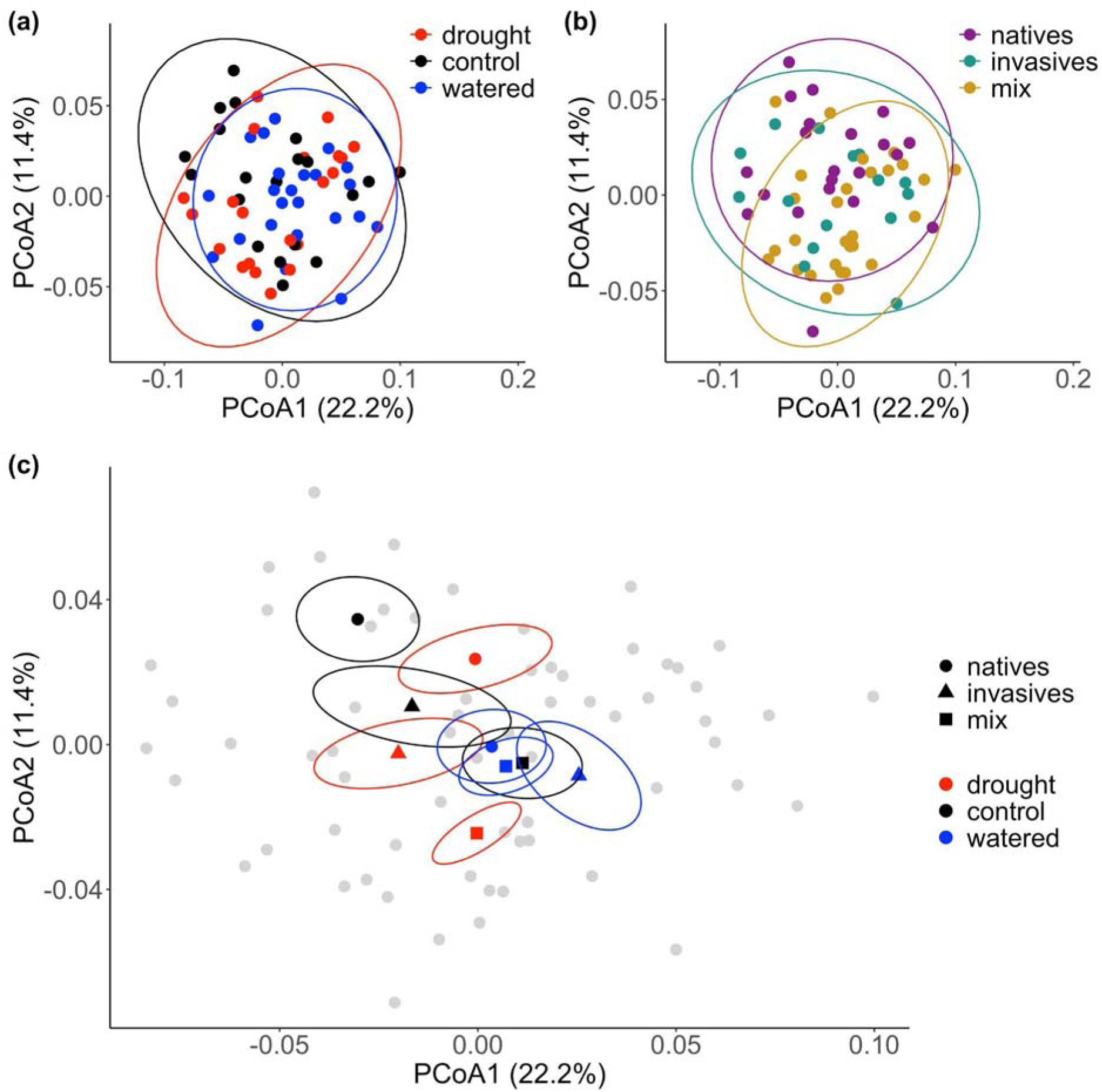
Ordination on Bray-Curtis dissimilarities of rhizosphere function from genes that appeared more than 3 times in at least 5% of the samples. (a) Principal coordinate analysis of functional beta diversity across watering treatments (control plots in black, watered plots in blue and drought plots in red). Functional beta diversity in the rhizosphere differed between watering treatments (p < 0.001). (b) Principal coordinate analysis of functional beta diversity across plant host treatments (native forbs in purple, invasive grasses in green, mixes in yellow). Functional beta diversity differed between plant host treatments (p = 0.044). (c) Centroids of microbial communities from each plant host (circles: native forbs, triangles: invasive grasses, squares: mixes) across watering treatments (drought: red, control: black, watered: blue). Note that scales between plots differ to allow for better visualization of patterns within each plot.

Due to variation in plant species composition within host plant treatments, we assessed whether overall host composition or presence of specific host species were drivers of beta diversity within treatment groups. Host plant composition was not significantly correlated with taxonomic beta diversity within invasive, native or native-invasive mixes (Figure S2; Mantel: *p* > 0.05), nor was the presence of individual host plant species (envfit for all species: *p* > 0.05). Similarly, overall host plant composition was not significantly correlated with functional beta diversity within invasive, natives or native-invasive mixes (Mantel: *p* > 0.05). While the presence of individual plant species was not a significant driver of the functional beta diversity of invasives or natives (envfit for all species: *p* > 0.05), the presence of two invasive grasses were significantly correlated with functional beta diversity in native-invasive mixes (envfit: *Elymus caput-medusae*: *p* = 0.039, *Festuca perennis*: *p* = 0.039).

### Invasive grasses interact with watering treatments to shift function and taxonomy in the native rhizosphere

While we did not find evidence supporting our hypothesis that the interaction between watering treatments and invasion were stronger than individual effects when investigating shifts in taxonomic and functional beta diversity, we instead found evidence supporting this hypothesis when assessing the differential abundance of gene function and taxonomy.

Using DESeq2, we identified 44 COG functions that significantly varied in abundance between treatments. We found no evidence that drought or watering alone shifts function in the native rhizosphere, nor that competition with natives shifts the function in the invasive rhizosphere. Instead, all differences in rhizosphere functions were found when comparing the native forbs growing alone to native forbs growing in competition with invasive grasses.

Analysis of the summed log-fold changes of these functions to the broader COG category level, revealed 16 COG categories (out of 25 total possible) that significantly varied between native rhizospheres and native-invasive mixed rhizospheres. Under control conditions, native-invasive mixes saw changes in relative abundance of four functional categories (FC) (Figure 3d). Specifically, we saw one function increase, COG3746 (Phosphate-selective porin OprP; FC = inorganic ion transport and metabolism), and observed a strong decrease in COG3592 (Uncharacterized Fe-S cluster protein Yjdl; FC = function unknown). Mixed communities under drought, however, saw significant changes in all 16 of these FCs compared to native communities in the control (Figure 3a), with two FCs showing significant increases and 13 FCs showing significant decreases. The one remaining FC (amino acid transport and metabolism) had a summed log-fold change close to 0. In particular, we found that FCs associated with posttranslational modification saw the largest increases in mixed communities under drought, while transcription and signal transduction mechanisms saw the largest decreases. In terms of specific COG functions, we saw the largest increases related to COG3158 (Potassium uptake protein Kup; FC = inorganic ion transport), COG1644 (Sigma 54 homolog RpoN; FC = transcription), and COG0339 (Zn-dependent oligopeptidase, M3 family Dcp; FC = post translational modification). While many functions decreased, we highlight decreases related to COG1276 (Putative copper export protein PcoD; FC = inorganic ion transport and metabolism) and COG3861 (Stress related protein YsnF; FC = function unknown).

**Figure 3.**
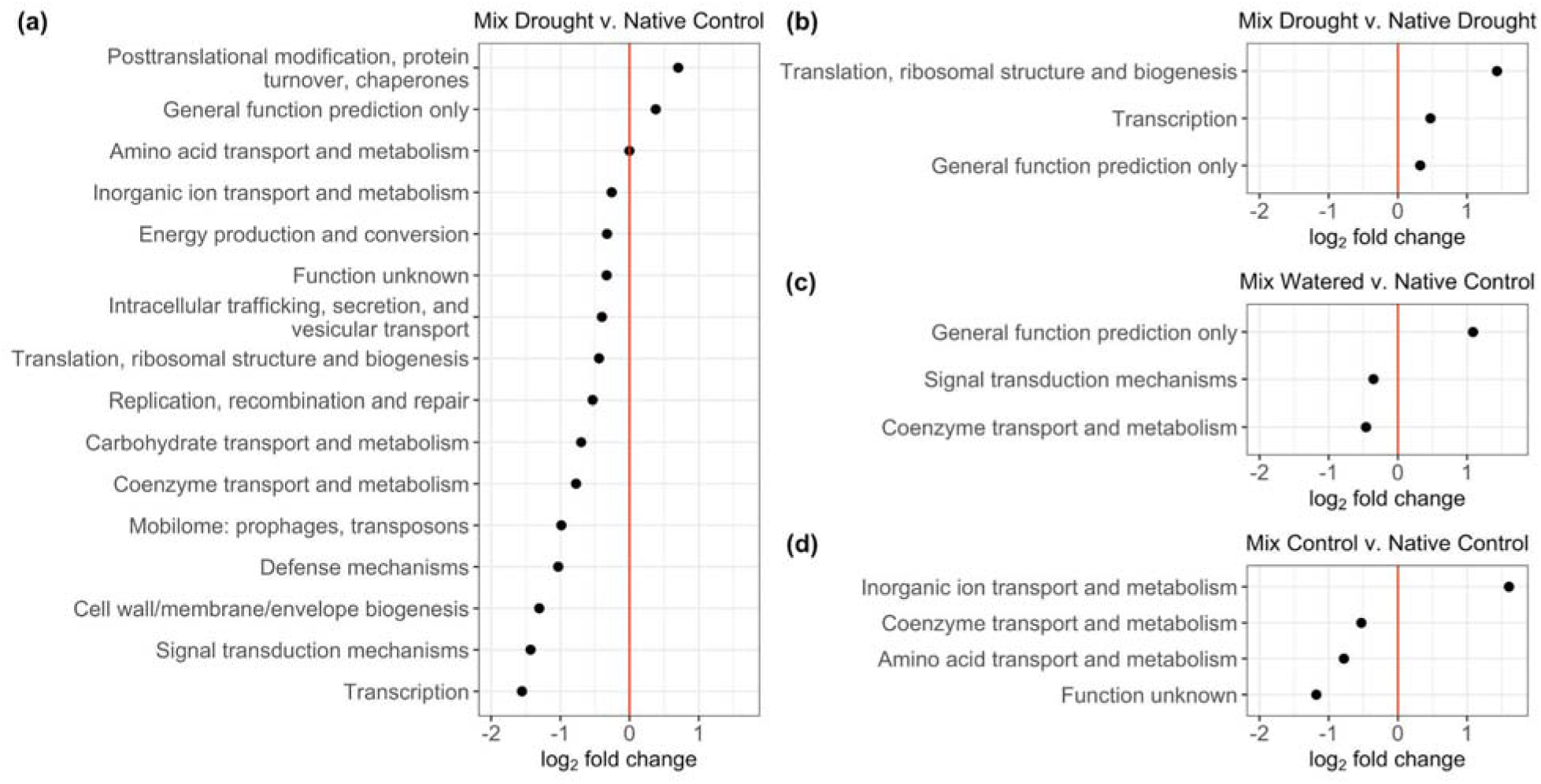
Log2 fold changes of COG gene categories between (a) mixes in drought and natives in control treatment; (b) mixes in drought and natives in drought treatment; (c) mixes in watering and natives in control treatment; (d) mixes in control and natives in control treatment. Values greater than zero indicate higher abundance in mixes, values less than zero indicate higher abundance in natives.

We also identified 3 FCs that significantly varied in mixed communities under drought compared to native communities under drought (Figure 3b; largest increase in translation) and 3 FCs that significantly varied between natives under control and native-invasive mixes under watering (Figure 3c). In terms of specific COG functions, native-invasive watered plots saw an increase in the COG3401 (Fibronectin type III domain FN3; FC = general function prediction only) and relatively small decreases in COG0161 (Adenosylmethionine-8-amino-7-oxononanoate aminotransferase BioA; FC = coenzyme transport and metabolism) and COG2205 (K+ sensing histidine kinase KdpD; FC = signal transduction mechanisms) (Figure S3).

Using DESeq2 we also identified 41 genera across 30 families that significantly varied in abundance between treatments with results largely mirroring functional results. That is, the majority of differences were observed between native-invasive mixes in drought and natives in control. Analysis of family-level summed log-fold changes identified 22 families that differed between mixes in drought and natives in control (Figure 4a), 5 families that differed between mixes in watered treatments and natives in controls (Figure 4b), and 16 families differed between mixes in controls and natives in controls (Figure 4c). The majority of these families belong to the phyla Actinobacteria (10 families) and Proteobacteria (10 families), with the majority of Actinobacteria families decreasing (8 families) and the majority of Proteobacteria families increasing during competition (10 families), with the strongest shifts under drought. Of the many families that saw changes in abundance, Sandaracinaceae (Proteobacteria), Hungateiclostridiaceae (Firmicutes), and Burkholderiales (Proteobacteria) all showed the largest increases under invasion and drought. Contrary to functional results, no differences were found between natives under drought and mixes under drought.

**Figure 4.**
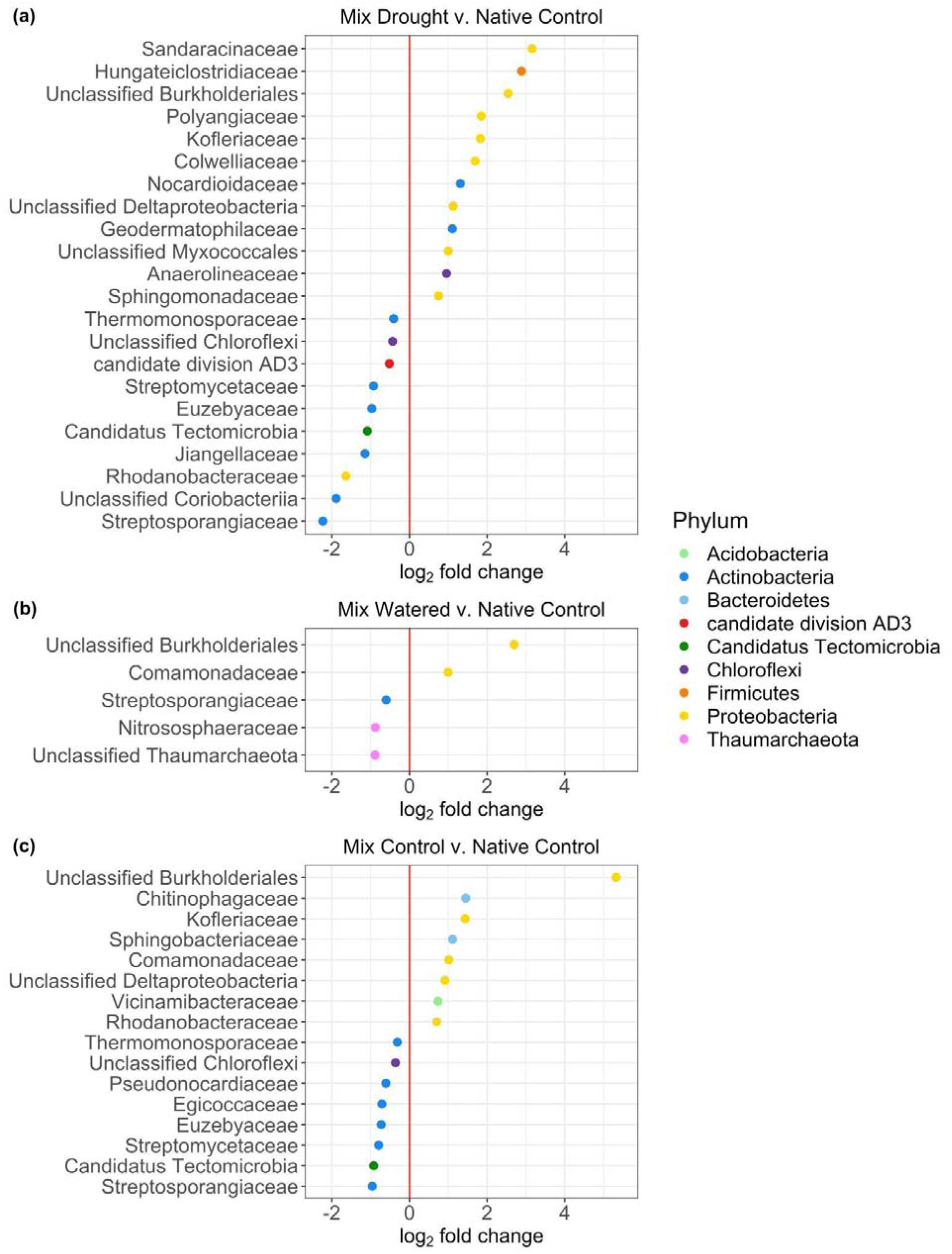
Log2 fold changes of taxonomic families between (a) mixes in drought and natives in control treatment; (a) mixes in watering and natives in control treatment; (c) mixes in control and natives in control treatment. Values greater than zero indicate higher abundance in mixes, values less than zero indicate higher abundance in natives.

### Under drought, the majority of the microbial community is assembled from invasive grasses

We assessed the contribution of native forbs and invasive grasses to microbial families in native-invasive subplots using SourceTracker2. We hypothesized that invasives would source a majority of the community during drought, but that the microbial community would be equally sourced in watered plots. Instead, we observed that across all mixed subplots, invasives were the main contributors to the communities, sourcing consistently over half of community membership (Figure 5). This was exacerbated during drought where invasives sourced on average 75.6% (± se: 6.89%) of the microbial community, a significantly higher proportion than was sourced by natives (mean ± se: 23.7% ± 6.87%; *p* < 0.001). Invasive and native contributions were also significantly different in control treatments (*p* = 0.019) and in watered plots (*p* = 0.015), with the contribution of the community sourced by natives highest during control treatments (native mean ± se: 35.6% ± 7.66%). Consistent with the overall community sourcing patterns, we found that differentially abundant families were on average sourced more by invasives and that this was heightened during drought (Figure 6).

**Figure 5.**
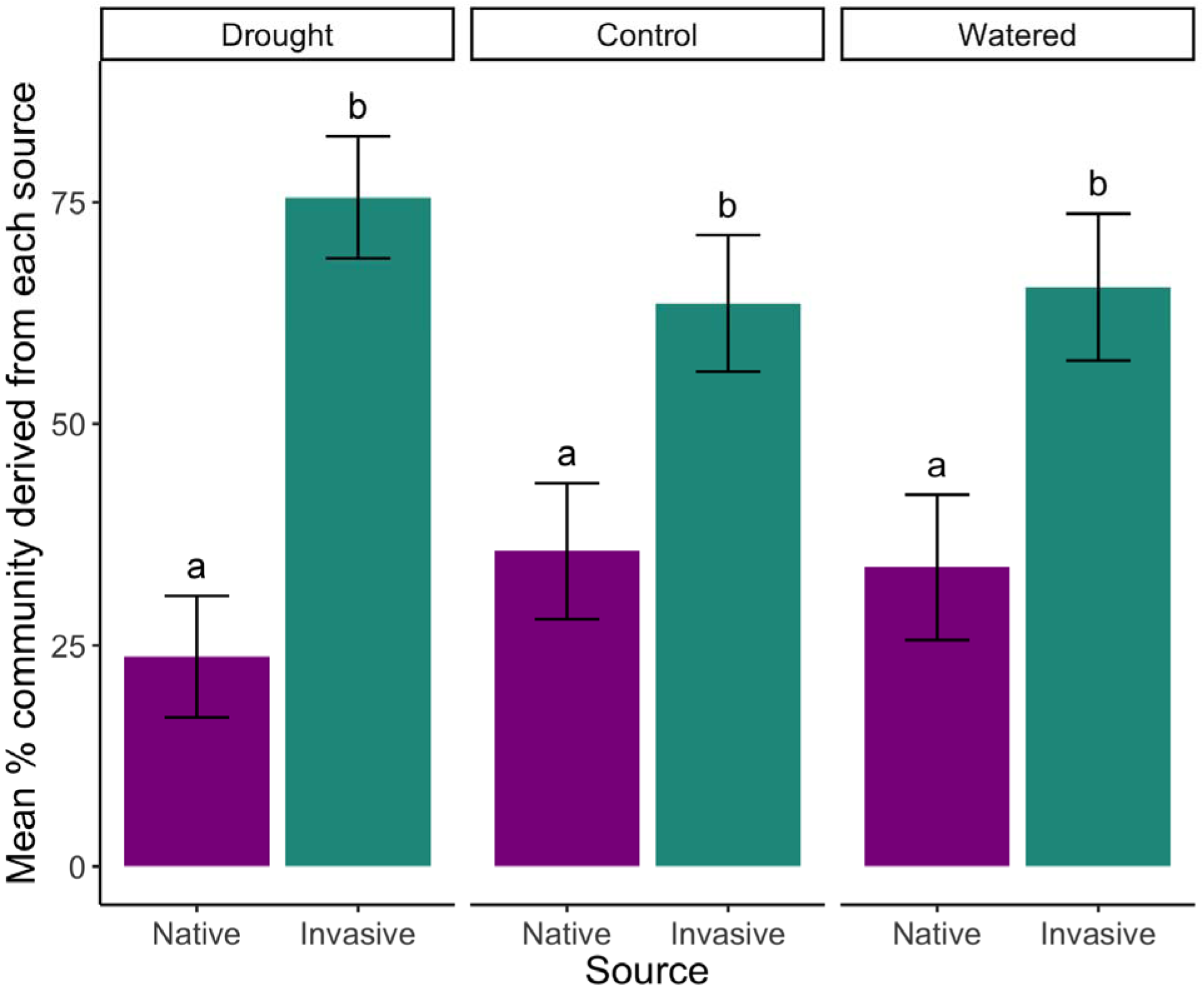
Invasives source more taxonomic families during drought. SourceTracker2 was used to predict whether genes associated with particular taxonomic families in treatments originated from invasive grasses (dark green), or native forbs (purple). Here we show the proportion of the community predicted to assemble from each source for each watering treatment. Comparisons that are significantly different from each other are notated by different letters (e.g., a vs. b). Error bars represent standard error.

**Figure 6.**
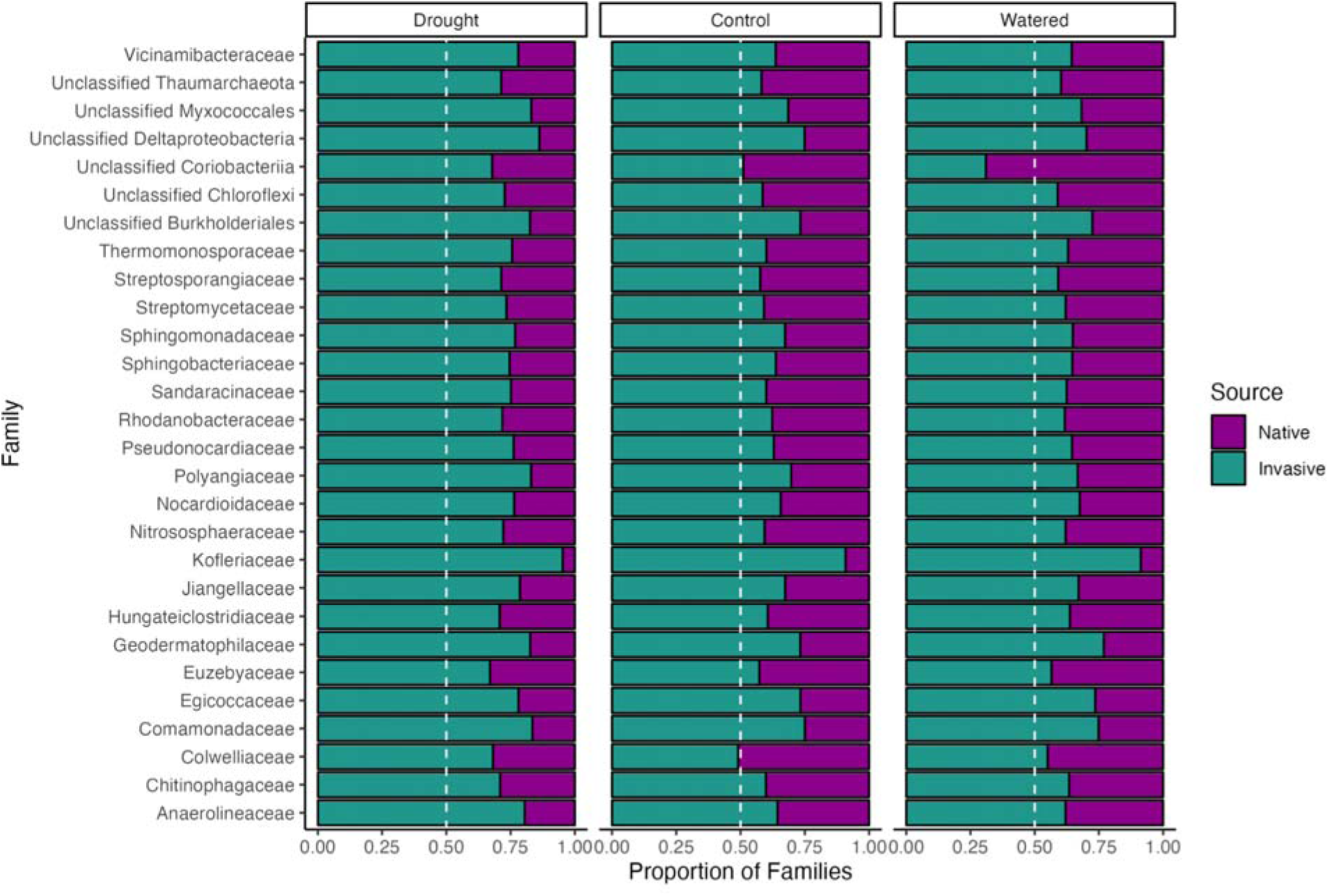
ST Families. Bacterial families containing significant differentially abundant genera are more sourced from invasives during drought. Proportion of families predicted to colonize from each source for each taxonomic family found to have differentially abundant genera between treatments. SourceTracker2 was used to predict whether genes associated with particular bacterial families in treatments originated from invasive grasses (dark green), or native forbs (purple). A dashed line is used to designate a hypothesis of equal contribution from both sources (i.e., 50% proportion).

We also used SourceTracker2 to investigate the origin of COG functions in native-invasive subplots. Again, invasives contributed significantly more to the functional potential of the community sourcing slightly more functions than natives (Figure S4). This was highest during drought where invasives sourced on average 50.9% (± se: 1.10%) of the functional potential of the community, a significant, but only slightly higher proportion than is sourced by natives (mean ± se: 44.9% ± 0.97%; *p* = 0.003). Invasive and native contributions were also significantly different in control treatments (*p* = 0.009) and in the watered treatments (*p* = 0.012), with the contribution of the community sourced by natives again highest during control treatments (native mean ± se: 46.3% ± 0.57%). Similar to the overall functional sourcing patterns, we found that differentially abundant COG functions were sourced fairly equally across treatments, with invasives providing a slightly higher proportion of the functional potential overall (Figure S5). One notable exception to this trend is the putative copper export protein (PcoD) function, which was sourced in its entirety by natives, regardless of watering treatment.

## Discussion

Our study adds to growing evidence that invasive species may affect the ability of native communities to cope with climate change and highlights that altering key functions and taxa in the rhizosphere microbiome is one pathway through which this may occur. While we did not find evidence for interactive effects of rainfall and plant host on microbial beta diversity, we found that beta diversity varied in response to the main effects of watering and host treatments and further, that drought combined with invasive competition led to shifts in abundance of more functions and families than invasive competition or drought alone. Thus, while overall community composition was not sensitive to interactive effects of invasive competition and watering, interacting global change factors have the potential to alter numerous specific functional pathways in native rhizosphere communities. Conversely, competition with native forbs did not appear to shift the abundance of specific functions or microbial families within invasive grasses regardless of watering treatment, indicating that invasive grass microbiomes are less affected by both competition and climate change in this community. Further, invasive grasses seem to be responsible for sourcing a majority of microbial families and functions in native-invasive mixes regardless of watering treatment, but with the highest sourcing of microbial families under drought. This suggests that invasives are likely exerting stronger selection on the microbial rhizosphere community than natives overall and that invasive-associated microbes may be able to outcompete native-associated microbes even during drought.

### Competition with invasives shifts native microbial communities

Invasive grass competition changed the overall community similarity between native rhizospheres, and led to enrichment in the abundance of specific families and functions. In previous work utilizing the same set of species, we found that taxonomic beta diversity significantly differed between invasive grasses, native forbs, and native-invasive pairs using 16S rRNA gene amplicon sequencing (LaForgia et al., 2022; Rodríguez-Caballero et al., 2020). While we did not find that here, it is possible that intra-group interactions (i.e., interactions between native forbs or between invasive grasses) in the field mask differences between natives and invasives as plants compete for limited nutrients (Hawkins & Crawford, 2018). Despite this, our findings that invasive competition shifts native rhizosphere microbes is generally well-supported (Komatsu & Simms, 2019; LaForgia et al., 2022; Rodríguez-Caballero et al., 2020), and we speculate that these invasive-associated microbial shifts play a role in the direction and magnitude of native-invasive interactions as seen elsewhere (Emam et al., 2014; Klironomos, 2002). The sole function that increased in abundance under invasive competition was OprP. OprP is expressed during phosphate starvation, has an affinity for orthophosphate (the preferred variety uptaken by plants), and genes with this annotation have been previously reported in the barley rhizosphere (Chhabra et al., 2013; Siehnel et al., 1988, 1992). While we do not know for certain that genes with this annotation perform the same function here, the enrichment of this function in native-invasive mixes may indicate competition for key nutrients by microbes during host competition.

Competition between natives and invasives also led to shifts in the abundance of numerous families including an enrichment of families in the phylum Proteobacteria, which tend to dominate serpentine soils (Mattarozzi et al., 2017) as well as grass rhizosphere microbiomes (B. K. Singh et al., 2007), and a decrease in families in Actinobacteria under competition. Both Actinobacteria and Proteobacteria are commonly enriched in serpentine soils, especially in the rhizospheres for species with high heavy metal tolerance (Abou-Shanab et al., 2009; Igwe et al., 2021; Mattarozzi et al., 2017; Navarro-Noya et al., 2010), thus it is interesting that these groups had divergent responses to our treatments. We speculate that, in this study, Proteobacteria are more closely associated with the fast-growing invasive grasses while Actinobacteria are more closely associated with heavy-metal tolerant natives. Thus, the differential responses observed may be due to adaptive differences of microbes within phyla - Actinobacteria tend to be slow-growing with low metabolic rates, while Proteobacteria are largely fast-growing with high metabolic rates (Mattarozzi et al., 2017).

Of particular note within Proteobacteria, we observed a greater than 5 log-fold increase in Burkholderiales in native-invasive mixes compared to native subplots. This corroborates our previous findings in which members of this family were predicted to be primarily sourced from invasive grasses and whose abundance was correlated with invasive biomass during competition (LaForgia et al., 2022). Members of the Burkholderiales are often beneficial endophytes of grasses and can provide antimicrobial, antifungal and nitrogen fixation functions (Banerjee et al., 2018; Depoorter et al., 2016; Dos Santos et al., 2012; Eberl & Vandamme, 2016; Shehata et al., 2016). Given the results of this and our previous work, we hypothesize that members of the Burkholderiales may play a direct or indirect role in aiding the competitive dominance of invasive grasses in California grassland ecosystems. Together, these findings suggest that invasive grasses may be shifting rhizosphere communities to their competitive benefit.

### Drought amplifies microbial effects of invasive competition

Although competition with invaders was associated with some shifts in the microbial community, the majority of changes occurred when natives competed with invasives under drought. Specifically, functions that were less abundant in native-invasive mixes under drought compared to natives in controls had annotations related to transport and metabolism, energy production, replication, mobilome, and defense. Many of these functions are related to secondary metabolism, which may provide a competitive advantage under local serpentine conditions (Haferburg et al., 2009; Porter et al., 2017), but also have a high energetic cost (Yan et al., 2018). Thus, they may be less advantageous under resource-limited conditions such as during drought when there is decreased microbial biomass and enzymatic activity (Baldrian et al., 2010; Sardans & Peñuelas, 2005). Alternatively, microbial taxa associated with these functions may be simply less abundant during drought, though this seems less probable given the observed function redundancy in these communities across watering treatments. Regardless, it is possible that a decrease in genes associated with secondary metabolism under drought may relate to a higher susceptibility of natives to invasive grass competition.

Genes with a higher abundance under drought in native-invasive mixes had putative functional annotations related largely to nutrient acquisition. For example, potassium is a critical macronutrient for microbes and their plant hosts (Pandey et al., 2020), and increased abundance of genes with predicted Kup, a potassium uptake protein, annotations may relate to increased competition for limited nutrients in native-invasive mixes during drought. Another putative function that increased, RpoN, can act as a regulator of nitrogen assimilation and virulence (Kullik et al., 1991), and in both capacities could provide competitive dominance for invasives through nutrient acquisition or microbial warfare with native-associated microbes, assuming that genes with this annotation perform similarly here. Further, Dcp, which also included genes that increased in mixes under drought, is reported to be involved in metabolism in cleaving a variety of bacterial peptides, and may relate to survival during carbon deficiency (Jasilionis & Kuisiene, 2015; Masuyer et al., 2016; Paschoalin et al., 2005; Vimr et al., 1983). Our findings also highlight that competing with invasive grasses under drought shifts rhizosphere function in a way that differs from the effect of drought on natives alone. Compared to natives exposed to drought, native-invasive mixes under drought displayed higher levels of genes associated with transcription and translation, possibly indicating a capacity for increased microbial activity in mixed subplots versus native subplots during drought. Thus, it is possible that invasive grasses under drought may be aided by their recruited microbiome at the expense of natives during competition.

Competition from invaders under drought was also associated with shifts in the abundance of 22 families (10 decreases, 12 increases). Largely, we saw an enrichment of families in the phylum Proteobacteria and Firmicutes, which have been previously reported to be abundant under drought conditions in grassland soils (Naylor & Coleman-Derr, 2017; Ochoa-Hueso et al., 2018; Xu et al., 2021; B. Zhang et al., 2020). While families in Actinobacteria saw a decrease under competition with invaders, especially under drought. Both Actinobacteria and Proteobacteria have been reported to be enriched under drought (Xu et al., 2021). Sandaracinaceae (Proteobacteria) and Hungateiclostridiaceae (Firmicutes), which were both sourced predominantly from grasses, especially under drought, include members that can degrade alternative carbon sources (Mohr et al., 2012; Rettenmaier et al., 2019, 2020; X. Zhang et al., 2018). This may be particularly beneficial under drought when there are reported changes in plant litter chemistry and carbon metabolism (Malik & Bouskill, 2022; Su et al., 2020). Lastly, Streptosporangiaceae (Actinobacteria) was consistently higher in native rhizospheres than mixes regardless of watering treatment, with the largest decline in abundance during competition with invasives under drought. Interestingly, despite its higher abundance in natives, this family was sourced more by invasives in mixes and this was exacerbated in drought. Members of Streptosporangiaceae have been isolated from soil and plant materials (Ara & Kudo, 2007; Janso & Carter, 2010; Kudo et al., 1993), and are thought to harbor novel and interesting antibiotic compounds (Lazzarini et al., 2000). These results, combined with the fact that invasive grasses sourced a higher proportion of microbial families under drought, underline the importance of investigating plant-microbe interactions across environmental gradients and add to a growing body of literature suggesting negative synergistic effects of interacting global change drivers.

### Effects of invasive competition under increased watering

Invasive grass competition under well-watered conditions drove substantially fewer changes in the microbial community compared to competition under drought, with native-invasive mixed subplots having less abundance of genes with predicted annotations related to signaling and metabolism, including BioA and KdpD. BioA is required for biotin synthesis, which is an important cofactor of carboxylases for rhizobia metabolism and root colonization (Heinz & Streit, 2003; Streit et al., 1996); BioA has also been predicted to confer copper resistance in *Burkholderia* (Higgins et al., 2020). KdpD is an osmosensitive sensor kinase related to potassium homeostasis (Lipa & Janczarek, 2020), which can activate under low potassium conditions and has been previously reported in rhizobia (Prell et al., 2012; Sablok et al., 2017). Potassium has been suggested to be a key signal of environmental cues as well as plant-microbe interactions and microbial virulence (Feng et al., 2022; Freeman et al., 2013; Tan, 2021). Given that both BioA and KdpD have been linked to stress response and documented roles in plant-microbe interactions, assuming they have similar functional roles here, they may be local adaptations by microbes to survival in serpentine soil, or possibly play roles in regulating native forb-microbe associations. Moreover, no families differentially varied between watered native-invasive mixes and native control subplots. High water levels may therefore ameliorate competition for below ground resources between native forbs and invasive grasses as competition for light aboveground intensifies (Coleman & Levine, 2006). In another study, the presence of invasive grasses dampened the beneficial effects of watering rather than exacerbating any negative effects (LaForgia et al., 2020). Thus, drought is likely to be a stronger driver of invasive-mediated microbiome shifts under future climates than high rainfall.

### Invasives drive community assembly dynamics during competition

Contrary to our hypothesis of equal contribution to the microbiome from natives and invaders, we observed that invaders sourced an overall higher proportion of the taxonomic families and functional potential in the rhizosphere microbiome. This contrasts with previous results from 16S rRNA gene amplicon work from a well-watered greenhouse experiment which found equal contribution by natives and invasives in native-forb pairs, despite higher biomass in grasses (LaForgia et al., 2022). While there are several possible explanations for this discrepancy, including methodological differences, one intriguing hypothesis is that the inclusion of multiple invasive species may result in a synergistic or additive competitive dominance over native rhizospheres (Kuebbing et al., 2013). While the overall taxonomic community was sourced predominantly by invasives, with an enrichment in drought, some taxonomic families (e.g. unclassified Coriobacteriia [Actinobacteria]) show a more complex pattern, including relatively higher sourcing from natives during well-watered treatments compared to control and drought treatments. While our taxonomic results suggest clear shifting due to selection by drought and/or invasives, the more subtle differences in functional sourcing suggest that there is likely selective pressure for community functional redundancy. This decoupling of taxonomic and functional results has been reported previously with taxonomic composition and functional potential being shaped by different abiotic and biotic factors (Louca et al., 2018).

The dominance of invasives as the source of microbes and functions regardless of watering treatment may not be surprising as invasive grasses account for 71% of plant cover and are the largest biomass contributor at these sites (LaForgia, unpublished data). However, the fact that microbial sourcing was unrelated to plant biomass in our previous work (LaForgia et al., 2022) suggests that microbial community assembly is likely to be more complex beyond host relative abundance. Further, invasive plants are known to modify the rhizosphere microbiome to negatively impact native-microbe interactions, as well as modify the soil chemistry to impact functional activity, with the intensity of these impacts increasing with invasive density (Fahey et al., 2020; H. Y. Zhang et al., 2020). In spite of invasive dominance, in mixed subplots we do see one function with differential abundance that is consistently sourced solely by native-associated microbes. That function, PcoD, encodes an internal membrane protein pump that works in copper translocation and is required for copper resistance (Rensing & Grass, 2003; Solioz, 2019). Copper is one of several heavy metals prevalent in serpentine soils that both microbes and plants need to evolve tolerance of to thrive (Fernandez et al., 1999; Llugany et al., 2003). The sourcing of this function highlights the importance of native plants for sustaining microbial functionality in grassland ecosystems given their evolution with local microbes. As climate change and invasion lead to shifts in the microbiome, we may start to see the loss of beneficial microbes as other taxa with the same functional potential are selected for or are brought in by invaders. These microbes may outcompete native-associated microbes and may not provide the same benefits to native plants that native-associated microbes would.

### Caveats

While metagenomics allows us to move beyond high throughput amplicon sequencing to ask questions about the functional potential of a community, it does not provide information about which genes are being actively expressed. Future work should use metatranscriptomics and/or metaproteomics to directly profile how invasion and climate change may interact to shape functional activity in the rhizosphere microbiome. Further, due to volunteer species, as well as species that were seeded but did not take, we had variable plant community composition within treatments and were only able to sequence 66 of 150 total samples, thus lowering replication and the power to identify differences. While we found that host plant composition was not a significant driver within treatments here, we know from other work the importance of host plant identity in shaping the microbial community (Aleklett et al., 2015). It is also possible that sampling root endospheres directly instead of the rhizosphere would have better let us detect host-specific signatures. However, we chose to sample the rhizosphere as that is where microbial crosstalk is likely occurring between competing microbes associated with invasive and natives (Sindhu et al., 2017) and is likely best to detect effects of competition. Regardless, our results are in line with other work showing that taxonomic beta diversity of the rhizosphere microbiome is heavily affected by drought (Ochoa-Hueso et al., 2018) and less so by plant species composition (Sayer et al., 2017), including invasion (Fahey et al., 2020). Similarly, the response of taxonomy to climate (Chen et al., 2022) and to invasion (Gibbons et al., 2017) has been shown to be stronger than functional response, likely due to functional redundancy across taxa.

## Conclusions

To summarize, we found that both drought and competition with invasives shift native microbial communities, and that the combined effects of drought and competition, while not observed at a community level, were seen when looking at specific functions and families. A majority of shifts were in functions associated with stress, starvation and survival in minimal nutrient conditions and include functions involved with efflux and transport of heavy metals. We saw increased abundance of Proteobacteria and decreased abundance of Actinobacteria during competition, with the strongest shifts during drought. Of the families that shifted of particular note is the invasive grass-associated Burkholderiales, which in combination with our previous work, seems to possibly be a key player in assisting invasives with their competitive dominance. Further community composition investigations revealed that invasives are the dominant source of functional potential and taxonomic identity in the microbial communities of invaded subplots regardless of rainfall variability. Altogether our results support growing literature suggesting that invasives may shift rhizosphere microbial communities for their benefit, possibly at the expense of natives and native-associated microbial communities.

## Funding Sources

This work was supported in part by grants from the UC Davis Center for Population Biology Collaborative Project research grant to CLE & MLL, UC Davis Center for Population Biology student grant to CLE, SIGMA XI Grant-In-Aid of Research to CLE, and UC Davis Center for Population Biology Hardman Foundation Research Award to MLL.

## Acknowledgments

We thank Cathy Koehler and Paul Aigner for their help setting up the watering experiment as well as Sonia L. Ghose and Andrew Latimer for their help with field sampling. We also thank members of the Hallett Lab, Gremer Lab, and Stajich Lab for their insightful feedback and valuable comments. Finally, we would like to thank Susan P. Harrison and Jonathan A. Eisen for useful conversations and suggestions related to this work.

## Notes

https://www.ncbi.nlm.nih.gov/bioproject/PRJNA925931

https://github.com/marinalaforgia/McL_Metagenomics

### Competing Interest Statement

The authors have declared no competing interest.

