## Supplementary Material for "Invasive plant species interact with drought to shift key functions and families in the native rhizosphere"

### S. Figure 1. Taxa PCoA

Ordinations on Bray-Curtis distances of rhizosphere taxonomic beta diversity from genes that appeared more than 3 times in at least 5% of the samples. (a) Principal coordinate analysis of taxonomic beta diversity across watering treatments (controls in black, watered in blue and drought in red). Taxonomic beta diversity in the rhizosphere differed between watering treatments (p < 0.001). (b) Taxonomic beta diversity also varied significantly across plant host treatments (p = 0.011); native forbs in purple, invasive grasses in green, mixes in yellow). (c) Centroids of taxonomic communities from each plant host (circles: native forbs, triangles: invasive grasses, squares: mixes) across watering treatments. Note that scales between plots differ to allow for better visualization of patterns within each plot.
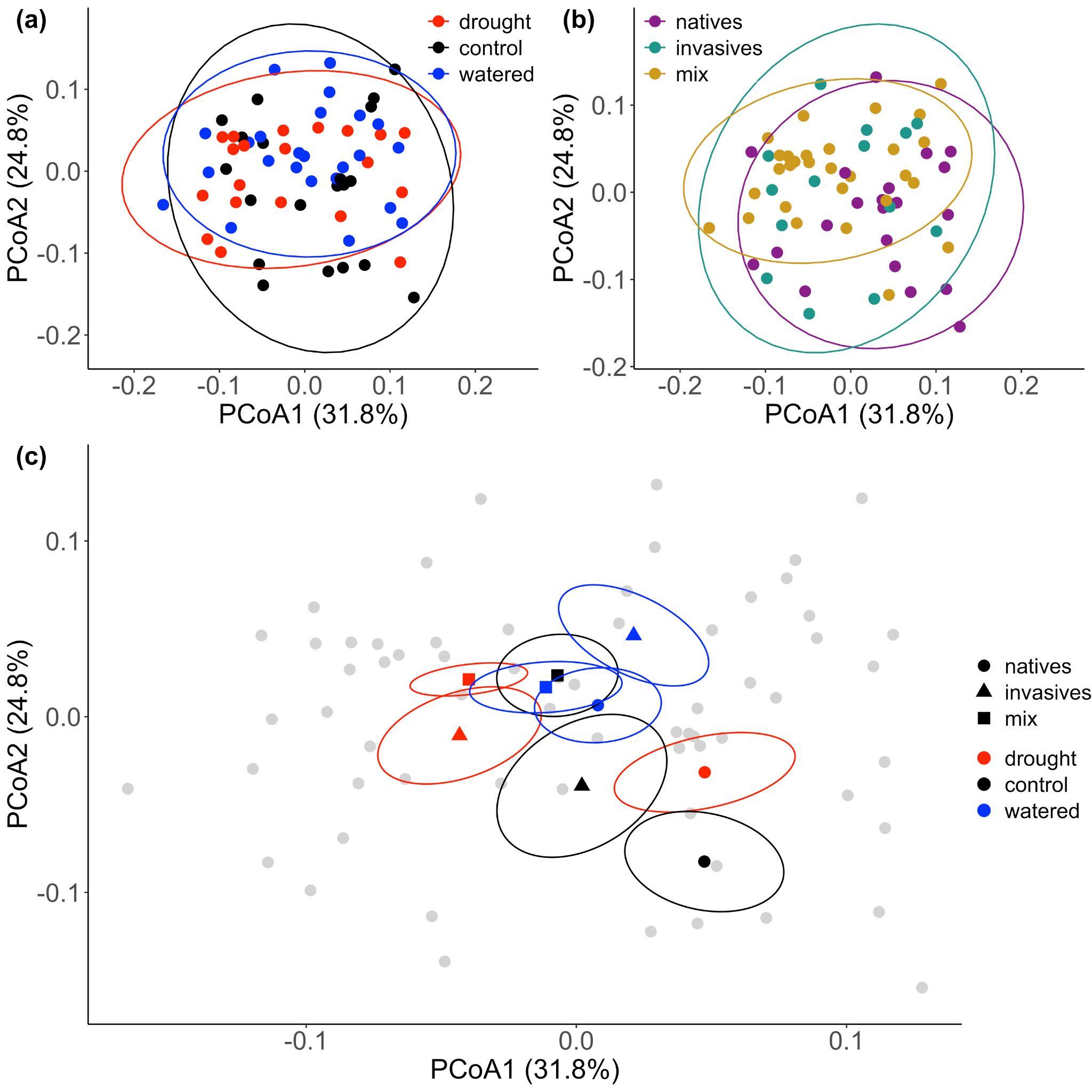

### S. Figure 2. Host PCoA

Host plant composition contribution to principal coordinate analysis of taxonomic beta diversity within invasives (a), natives (c) and native-invasive mixes (e), and functional beta diversity within invasives (b), natives (d) and native-invasive mixes (f). (a) Neither overall host plant composition or individual host plant species presence were significantly correlated with the taxonomic beta diversity of invasives (Mantel: *p* = 0.801; envfit for all species: *p* > 0.05). (b) Similarly, neither overall host plant composition nor individual host plant species presence were significantly correlated with functional beta diversity of invasives (Mantel: *p* = 0.951; envfit for all species: *p* > 0.05). (c) While taxonomic beta diversity was significant (permanova, *p* = 0.04) between native mixes, overall host plant composition was not (Mantel: *p* = 0.386) nor was the presence of individual host plant species significant (envfit for all species: *p* > 0.05). (d) For functional beta diversity, overall host plant composition and individual species presence were not significant between native mixes (permanova: *p* = 0.089; Mantel: *p* = 0.338, envfit for all species: *p* > 0.05). (e) Similarly, taxonomic beta diversity was marginally significant (permanova: *p* = 0.068) between native-invasive mixes, while overall host plant composition and individual species presence were not (Mantel: *p* = 0.148; envfit for all species: *p* > 0.05). (f) Functional beta diversity however was significant between native-invasive mixes (permanova: *p* = 0.012). While overall host composition was not significant (Mantel: *p* = 0.158), the presence of two invasive grasses were significant (envfit: *Elymus caput-medusae*: *p* = 0.039, *Festuca perennis*: *p* = 0.039). Black arrows and text are used to indicate significantly correlated host plant species.

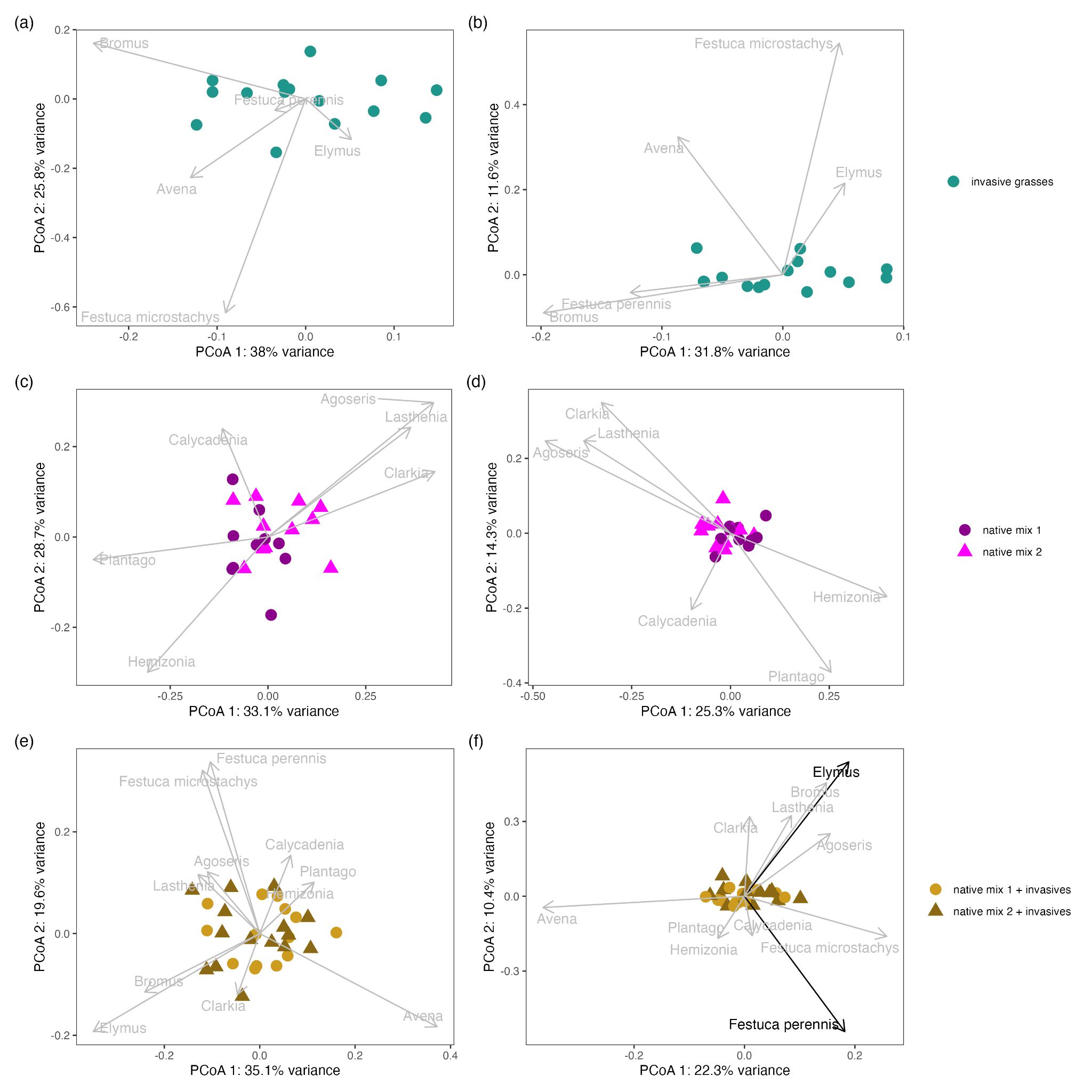

### S. Figure 3. DESeq2 COG20 functions

Log2 fold changes of COG20 functions colored by COG20 category. (a) Mixes in drought and natives in control treatment; (b) mixes in watering and natives in control treatment; (c) mixes in drought and natives in drought; (d) mixes in control and natives in control treatment. Values greater than zero indicate higher abundance in mixes, values less than zero indicate higher abundance in natives.

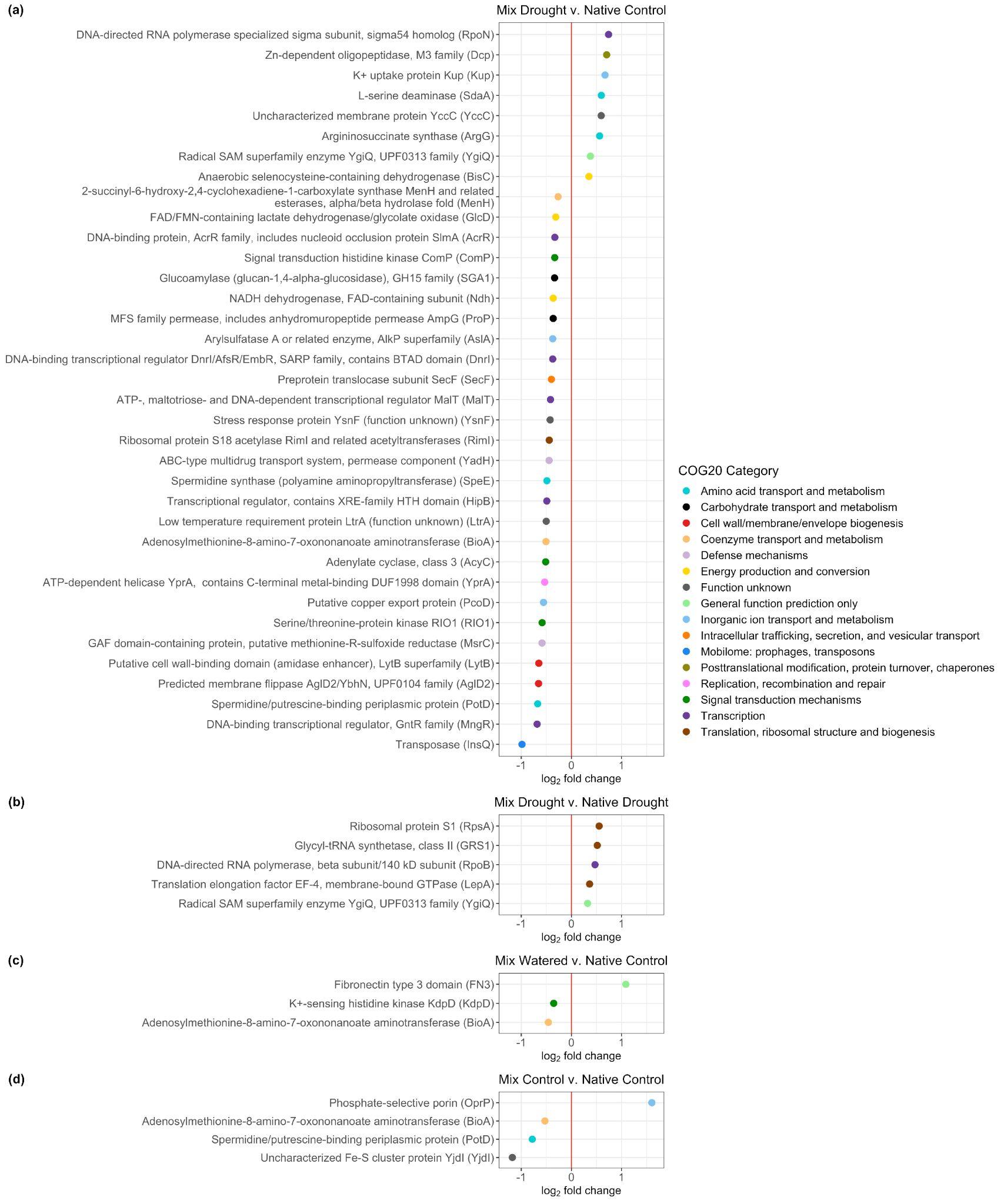

### S. Figure 4.

Invasives source slightly more COG functions during drought. SourceTracker2 was used to predict whether genes associated with particular COG functions in treatments originated from invasive grasses (dark green), or native forbs (purple). Comparisons that are significantly different from each other are notated by different letters (e.g., a vs. b). Error bars represent standard error.

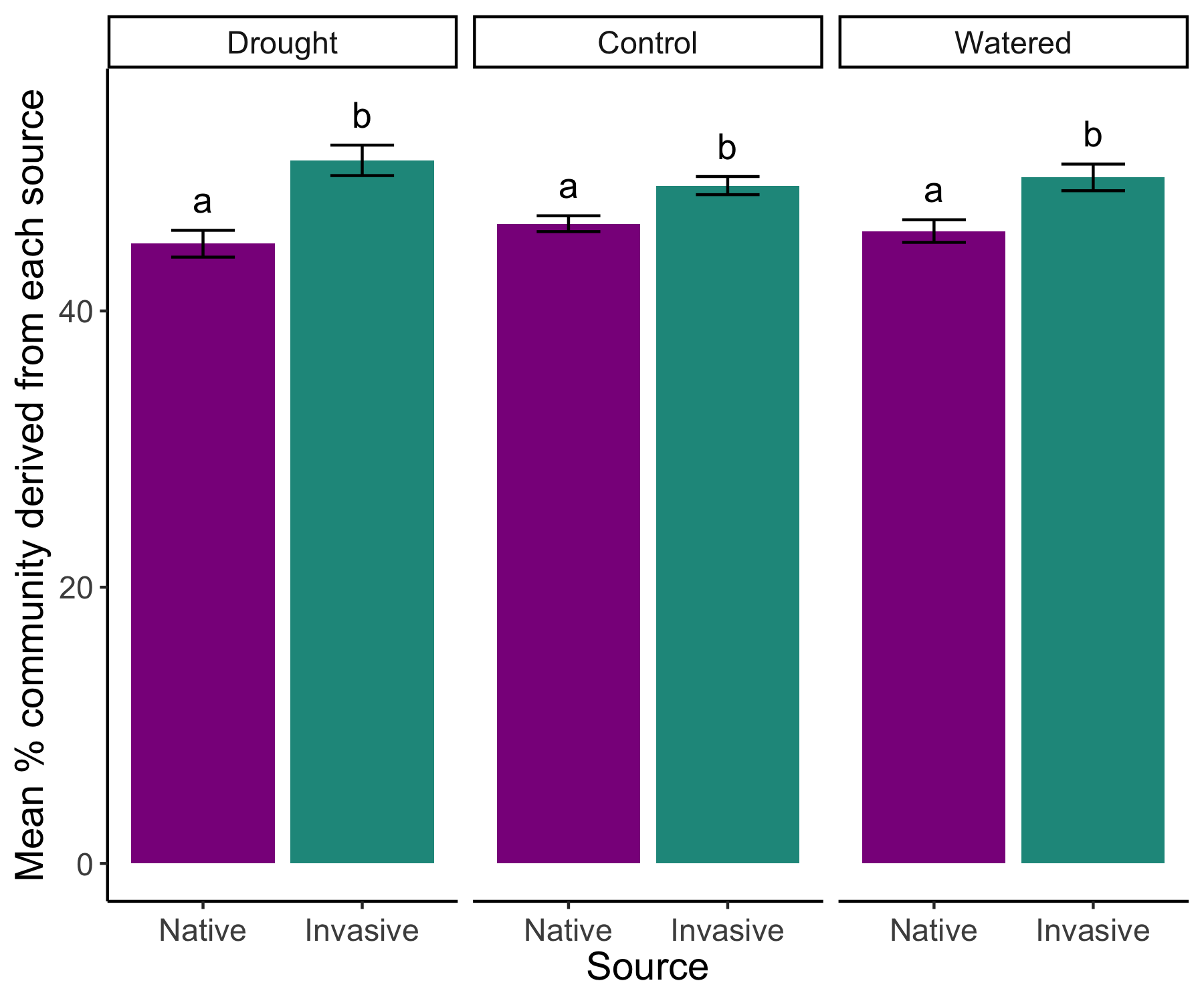

### S. Figure 5.

COG functions that are differentially abundant are sourced equally or slightly more by invasives, regardless of treatment. Proportion of families predicted to colonize from each source for each COG function linked to the significant COG functions that differed between treatments. SourceTracker2 was used to predict whether genes associated with particular COG functions in treatments originated from invasive grasses (dark green), or native forbs (purple). A dashed line is used to designate a hypothesis of equal contribution from both sources (i.e., 50% proportion).

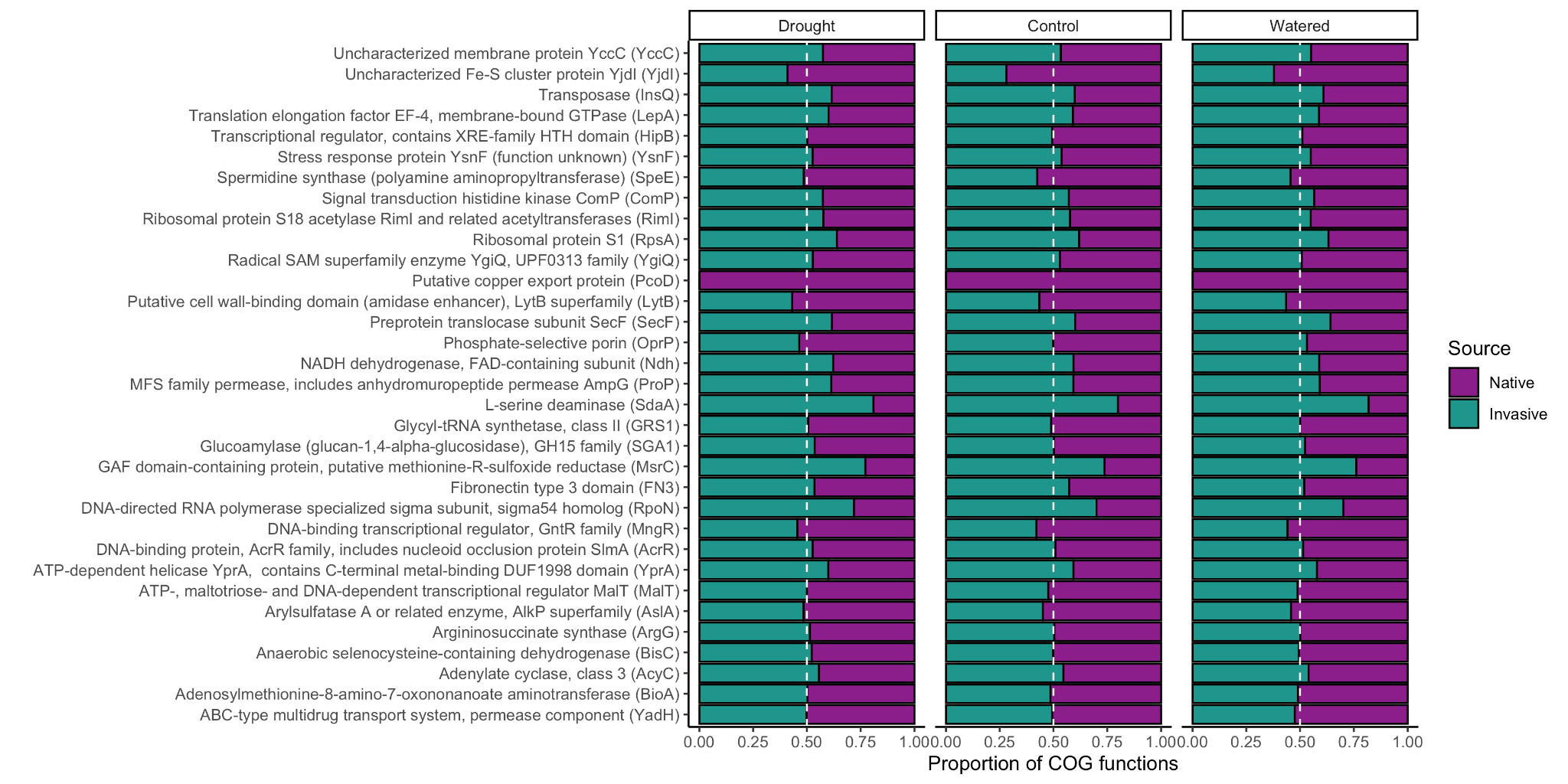

### S. Table 1.

Final plant species composition of each rhizosphere sample that was sequenced. A 0 indicates that species was not present within the core taken and a 1 indicates that species was present. We prioritized sequencing samples that included a majority of the original species seeded and did not have volunteer species that would shift the treatment functional group (e.g., we excluded native forb treatment subplots that had invasive grass volunteers). There were two instances in which a native grass (*Festuca microstachys*) appeared in mixed native-invasive subplots. This grass is in low abundance at the site and is much smaller than surrounding invasive grasses.

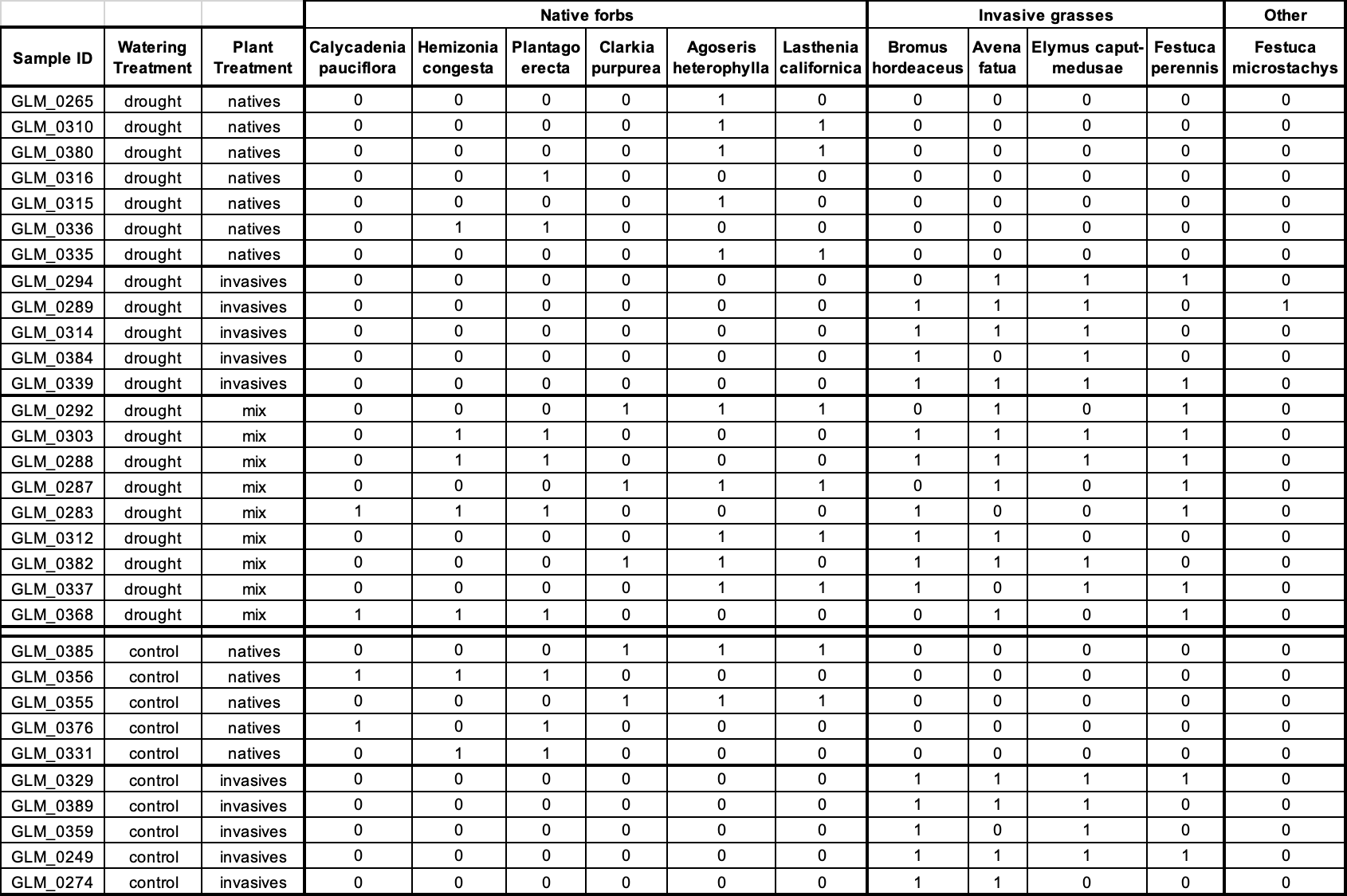

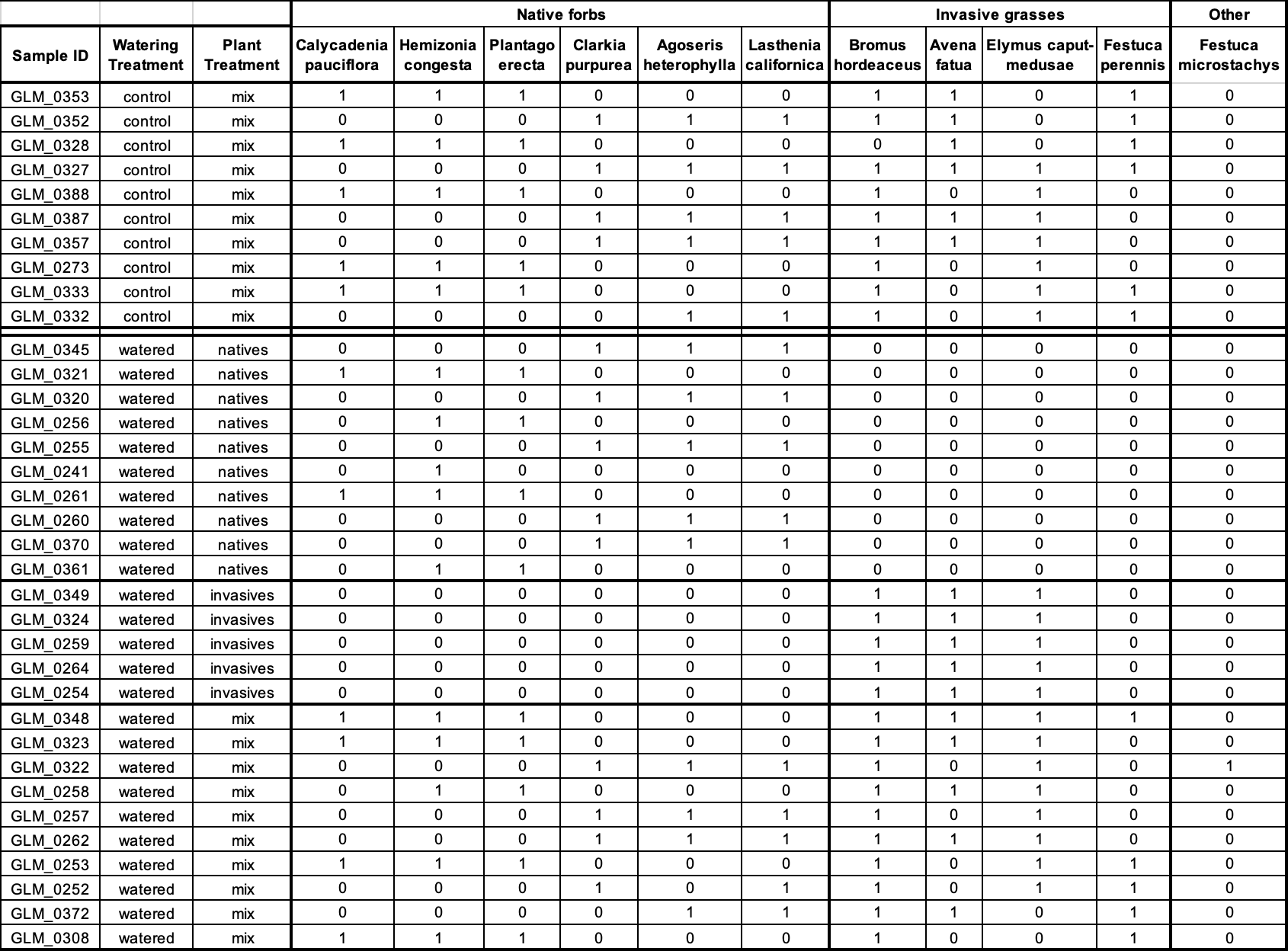

### S. Table 2.

Library sizes (total number of reads) and NCBI GenBank SRA accession numbers for each metagenomic sample in this work. Negative controls (GLM_0393, GLM_697) had an order of magnitude smaller library sizes than true experimental samples.

| **Sample** | **Type** | **Library Size** | **SRA accession no.** |
| --- | --- | --- | --- |
| GLM_0241 | Watered x mix | 8,151,242 | SRR23173716 |
| GLM_0249 | Control x invasives | 9,787,118 | SRR23173717 |
| GLM_0252 | Watered x mix | 13,523,892 | SRR23173718 |
| GLM_0253 | Watered x mix | 7,617,258 | SRR23173665 |
| GLM_0254 | Watered x invasives | 10,392,120 | SRR23173666 |
| GLM_0255 | Watered x mix | 8,803,852 | SRR23173667 |
| GLM_0256 | Watered x mix | 6,458,262 | SRR23173668 |
| GLM_0257 | Watered x mix | 11,083,658 | SRR23173669 |
| GLM_0258 | Watered x mix | 11,217,232 | SRR23173670 |
| GLM_0259 | Watered x invasives | 4,618,320 | SRR23173672 |
| GLM_0260 | Watered x mix | 1,913,950 | SRR23173673 |
| GLM_0261 | Watered x mix | 9,273,088 | SRR23173674 |
| GLM_0262 | Watered x mix | 15,920,680 | SRR23173675 |
| GLM_0264 | Watered x invasives | 6,531,154 | SRR23173676 |
| GLM_0265 | Drought x mix | 13,248,750 | SRR23173677 |
| GLM_0273 | Control x mix | 9,694,920 | SRR23173678 |
| GLM_0274 | Control x invasives | 9,209,114 | SRR23173679 |
| GLM_0283 | Drought x mix | 8,958,170 | SRR23173680 |
| GLM_0287 | Drought x mix | 10,100,112 | SRR23173681 |
| GLM_0288 | Drought x mix | 8,454,504 | SRR23173683 |
| GLM_0289 | Drought x invasives | 8,958,274 | SRR23173684 |
| GLM_0292 | Drought x mix | 12,012,520 | SRR23173685 |
| GLM_0294 | Drought x invasives | 11,982,616 | SRR23173686 |
| GLM_0303 | Drought x mix | 9,963,090 | SRR23173687 |
| GLM_0308 | Watered x mix | 7,585,572 | SRR23173688 |
| GLM_0310 | Drought x mix | 8,361,906 | SRR23173689 |
| GLM_0312 | Drought x mix | 9,133,838 | SRR23173690 |
| GLM_0314 | Drought x invasives | 11,741,128 | SRR23173691 |
| GLM_0315 | Drought x mix | 5,785,114 | SRR23173692 |
| GLM_0316 | Drought x mix | 6,791,226 | SRR23173694 |
| GLM_0320 | Watered x mix | 10,644,414 | SRR23173695 |
| GLM_0321 | Watered x mix | 16,911,776 | SRR23173696 |
| GLM_0322 | Watered x mix | 10,535,688 | SRR23173697 |
| GLM_0323 | Watered x mix | 8,590,358 | SRR23173698 |
| GLM_0324 | Watered x invasives | 10,138,428 | SRR23173699 |
| GLM_0327 | Control x mix | 7,402,208 | SRR23173700 |
| GLM_0328 | Control x mix | 7,065,688 | SRR23173701 |
| GLM_0329 | Control x invasives | 12,465,704 | SRR23173702 |
| GLM_0331 | Control x mix | 12,332,072 | SRR23173703 |
| GLM_0332 | Control x mix | 10,359,316 | SRR23173705 |
| GLM_0333 | Control x mix | 5,011,278 | SRR23173706 |
| GLM_0335 | Drought x mix | 9,328,720 | SRR23173707 |
| GLM_0336 | Drought x mix | 12,078,848 | SRR23173708 |
| GLM_0337 | Drought x mix | 9,745,972 | SRR23173709 |
| GLM_0339 | Drought x invasives | 8,737,910 | SRR23173710 |
| GLM_0345 | Watered x mix | 10,869,654 | SRR23173711 |
| GLM_0348 | Watered x mix | 8,959,182 | SRR23173713 |
| GLM_0349 | Watered x invasives | 8,447,728 | SRR23173719 |
| GLM_0352 | Control x mix | 6,572,610 | SRR23173720 |
| GLM_0353 | Control x mix | 8,499,492 | SRR23173722 |
| GLM_0355 | Control x mix | 7,548,688 | SRR23173723 |
| GLM_0356 | Control x mix | 8,361,988 | SRR23173724 |
| GLM_0357 | Control x mix | 5,137,396 | SRR23173725 |
| GLM_0359 | Control x invasives | 9,918,228 | SRR23173726 |
| GLM_0361 | Watered x mix | 10,308,956 | SRR23173727 |
| GLM_0368 | Drought x mix | 6,610,576 | SRR23173728 |
| GLM_0370 | Watered x mix | 10,617,230 | SRR23173729 |
| GLM_0372 | Watered x mix | 12,691,696 | SRR23173730 |
| GLM_0376 | Control x mix | 8,603,614 | SRR23173731 |
| GLM_0380 | Drought x mix | 9,139,006 | SRR23173712 |
| GLM_0382 | Drought x mix | 6,357,498 | SRR23173714 |
| GLM_0384 | Drought x invasives | 12,905,786 | SRR23173715 |
| GLM_0385 | Control x mix | 13,105,060 | SRR23173671 |
| GLM_0387 | Control x mix | 6,427,800 | SRR23173682 |
| GLM_0388 | Control x mix | 7,895,362 | SRR23173693 |
| GLM_0389 | Control x invasives | 5,481,064 | SRR23173704 |
| GLM_0393 | Negative control (sterile water) | 65,074 | SRR23173721 |
| GLM_0695 | Positive control (Zymo mock community) | 12,694,214 | SRR23173732 |
| GLM_0697 | Negative control (DNA extraction kit control) | 27,026 | SRR23173733 |

### S. Table 3.

Post-hoc contrasts of plant hosts and watering treatments for beta-diversity analyses of function and taxonomy. Bolded rows indicate Benjamini-Hochberg adjusted *p*-values less than 0.05. Differences in both functional and taxonomic beta diversity was apparent between native forbs and native-invasive mixes, and between all combinations of watering treatments. No differences were found between natives and invasives, or between invasives and mixes.

| **Beta-diversity type** | **Comparison** | **Adjusted *p*-value** |
| --- | --- | --- |
| function | natives v invasives | 0.369 |
| **function** | **natives v mixes** | **0.024** |
| function | invasives v mixes | 0.438 |
| **function** | **watered v drought** | **0.003** |
| **function** | **watered v control** | **0.014** |
| **function** | **drought v control** | **0.012** |
| taxonomy | natives v invasives | 0.156 |
| **taxonomy** | **natives v mixes** | **0.009** |
| taxonomy | invasives v mixes | 0.444 |
| **taxonomy** | **watered v drought** | **<0.001** |
| **taxonomy** | **watered v control** | **0.023** |
| **taxonomy** | **drought v control** | **0.012** |
